# Identification of a motif in TPX2 that regulates spindle architecture in *Xenopus* egg extracts

**DOI:** 10.1101/2024.02.10.579770

**Authors:** Guadalupe E. Pena, Xiao Zhou, Lauren Slevin, Christopher Brownlee, Rebecca Heald

**Affiliations:** Department of Molecular and Cell Biology, University of California, Berkeley, Berkeley, CA, USA; AbbVie, South San Francisco, CA, USA; Swedish Maternal and Fetal Specialty Center, Seattle, WA, USA; Deparment of Pharmacological Sciences, Stony Brook University, Stony Brook, NY, USA

## Abstract

A bipolar spindle composed of microtubules and many associated proteins functions to segregate chromosomes during cell division in all eukaryotes, yet spindle size and architecture varies dramatically across different species and cell types. Targeting protein for Xklp2 (TPX2) is one candidate factor for modulating spindle microtubule organization through its roles in branching microtubule nucleation, activation of the mitotic kinase Aurora A, and association with the kinesin-5 (Eg5) motor. Here we identify a conserved nuclear localization sequence (NLS) motif, ^123^KKLK^126^ in *X. laevis* TPX2, which regulates astral microtubule formation and spindle pole morphology in *Xenopus* egg extracts. Addition of recombinant TPX2 with this sequence mutated to AALA dramatically increased spontaneous formation of microtubule asters and recruitment of phosphorylated Aurora A, pericentrin, and Eg5 to meiotic spindle poles. We propose that TPX2 is a linchpin spindle assembly factor whose regulation contributes to the recruitment and activation of multiple microtubule polymerizing and organizing proteins, generating distinct spindle architectures.

## Introduction

During cell division, a dynamic, microtubule (MT)-based spindle forms to faithfully segregate duplicated chromosomes to daughter cells. Although the spindle performs this universal function in all eukaryotes, its architecture varies dramatically, both across species and in different cell types (Crowder et al., 2015). For example, during meiosis, a small, diamond-shaped spindle in the egg segregates half of the maternal genome to a small polar body. Following fertilization, the first mitotic division occurs and is accompanied by dramatic changes in spindle size and architecture, with long astral MTs extending outward from the spindle poles. These changes in spindle dimensions and architecture are likely to be critical for proper spindle function, but how they arise at the molecular level, remains unclear.

One candidate factor for modulating astral MT density and organization is the MT-associated protein TPX2. Originally identified as the spindle *t*argeting *<u>p</u>*rotein for the *Xenopus* kinesin motor *<u>X</u>*klp*<u>2</u>* (Wittmann et al., 2000), TPX2 is essential for spindle bipolarity, MT nucleation, stabilization and organization at spindle poles in multiple species and cell types (Brunet et al., 2004; Schatz et al., 2003; Tulu et al., 2006), and its overexpression is common in a number of cancers (Asteriti et al., 2010; Neumayer et al., 2014). TPX2 is one of several spindle assembly factors (SAFs) containing nuclear localization sequence (NLS) motifs that mediate binding and inhibition by importins, which is locally derepressed in the vicinity of mitotic chromatin by RanGTP (Gruss et al., 2001, 2002; Schatz et al., 2003). TPX2 drives MT nucleation at least in part by recruiting and activating Aurora A kinase, which phosphorylates targets such as NEDD1 to promote MT nucleation via γTuRC (Bayliss et al., 2003; Eyers et al., 2003; Garrett et al., 2002; Kufer et al., 2002; Pinyol et al., 2013; Sardon et al., 2008; Scrofani et al., 2015; Tsai et al., 2003). In addition, the C-terminal domain of TPX2 interacts with the homotetrameric plus end–directed motor Eg5, which crosslinks and slides antiparallel MTs outward (Eckerdt et al., 2008; Ma et al., 2011) and bundles parallel MTs at spindle poles (Helmke & Heald, 2014). TPX2 binding to Eg5 inhibits its ability to slide MTs *in vitro* (Gable et al., 2012; Ma et al., 2010, 2011). More recently, it was discovered that TPX2 plays a key role in branching MT nucleation off the sides of existing MTs together with augmin that recruits γTuRC and the MT polymerase XMAP215 (Kraus, Travis, et al., 2023; Petry et al., 2013; Song et al., 2018; Thawani et al., 2018; Travis et al., 2022). Both *in vitro* and in the presence of *Xenopus* egg cytoplasm, TPX2 is capable of undergoing liquid-liquid phase separation (LLPS), forming co-condensates with tubulin that stimulate branching MT nucleation (King & Petry, 2020). However, whether and how TPX2 is regulated to generate different spindle MT architectures and the underlying mechanisms are poorly understood.

*Xenopus* egg extracts provide an ideal system to study the role of TPX2 in defining spindle architecture since *in vitro* reactions combining cytostatic factor (CSF) metaphase-arrested extracts and sperm nuclei recapitulate meiotic spindle assembly and can be biochemically manipulated. Here we show that addition of recombinant *X. laevis* or *X. tropicalis* TPX2 in which a high affinity NLS motif, ^123^KKLK^126^ (Safari et al., 2021) is mutated to AALA caused dramatic changes in spindle architecture, MT polymerization, and astral MT growth, recruiting activated Aurora A, pericentrin, and Eg5 to spindle poles. *X. tropicalis* TPX2 possessed higher activity than the *X. laevis* protein, which was further enhanced by the NLS mutation. Interestingly, importin-α binding and recruitment to spindle poles occurred despite mutation of the NLS. We propose that TPX2-dependent association with multiple SAFs and importin-α promotes multivalent interactions that strongly enhance MT polymerization, branching, and bundling.

## Results and Discussion

### Mutating the 123-KKLK-126 motif *of X. laevis* and *X. tropicalis* TPX2 alters spindle pole morphology in *Xenopus* egg extracts

We showed previously that recombinant *X. laevis* and *X. tropicalis* TPX2 proteins produce distinct spindle architectures in egg extracts due to differences in TPX2 levels that affect spindle length and mid-zone organization, as well as a 7-aa deletion (Δ7) adjacent to the Eg5-binding domain in *X. tropicalis* TPX2 that was shown to stimulate astral MT growth and branching MT nucleation (Helmke & Heald, 2014). Through further mutational analysis of *X. laevis* TPX2, we identified another motif at 123-KKLK-126 that regulates MT polymerization in egg extract (Supplementary Figure 1). This NLS motif, termed NLS3, is conserved in mammals and contributes to the high affinity interaction between TPX2 and importin-α, which was shown to regulate TPX2 LLPS and co-condensation with tubulin (King & Petry, 2020; Safari et al., 2021). However, the function of this motif in regulating TPX2 interactions with other SAFs and MT architecture in the spindle was unknown.

To probe the role of NLS3 during spindle assembly, wildtype (123-KKLK-126) and NLS3 mutant (123-AALA-126) GFP-tagged *X. laevis* and *X. tropicalis* TPX2 proteins were expressed in *E.coli*, purified, and added to *X. laevis* egg extract spindle assembly reactions at 100 nM, similar to the endogenous concentration and below the 200 nM needed to significantly reduce spindle length (Helmke & Heald, 2014). Quantification of total GFP immunostaining on the spindle normalized to tubulin showed that NLS3 mutants of both species localized more strongly to spindles than wildtype proteins and altered spindle morphology, increasing MT density and frequently disrupting focusing of MTs to generate “hollow” spindle poles (Figure 1A and B). Additionally, mutant proteins induced ectopic MT aster formation (Figure 1C). The *X. tropicalis* wildtype and NLS3 mutant proteins exhibited stronger promoting of MT polymerization activity compared to the *X. laevis* versions, consistent with previous work (Helmke & Heald, 2014) and indicating an additive effect with the *X. tropicalis* 7-aa deletion. Thus, the NLS3 motif functions to regulate spindle architecture and MT aster formation and its mutation further increases the higher intrinsic MT nucleation activity of *X. tropicalis* TPX2.

**FIGURE 1:**
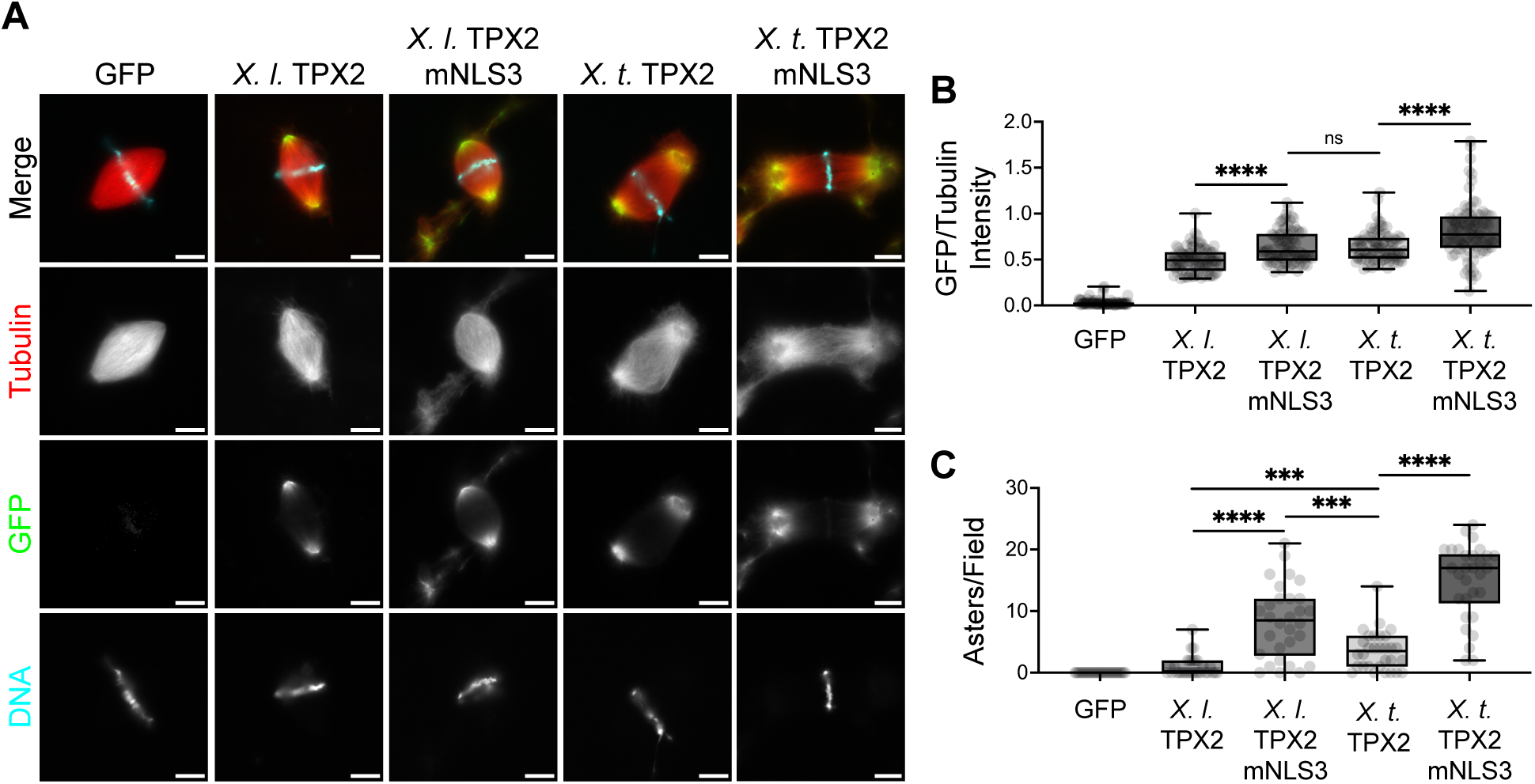
Addition of recombinant *X. laevis (X. l.)* and *X. tropicalis (X. t.)* mNLS3 TPX2 proteins modify spindle architecture in *Xenopus laevis* egg extract. (A) Immunostaining of spindle assembly reactions in the presence of 100 nM recombinant GFP or GFP-tagged TPX2 proteins. Spindle reactions immunostaining described in materials and methods. Scale bars = 10 µm. *X.t*. TPX2 mNLS3 image is cropped to show a larger field of view. (B) Ratio of fluorescence intensity of GFP/Tubulin on spindles, from n = 3 extracts, >25 spindles per biological replicate. For each TPX2 protein compared with GFP control **** = p<0.0001 from two-sided unpaired Welch’s *t* test. (C) Quantification of ectopic MT asters induced by recombinant TPX2 proteins. For each condition, the number of ectopic asters was counted in 10 microscope fields, n = 3 extracts. For each TPX2 protein compared with GFP control, **** = p< 0.0001 from unpaired Welch’s *t* test. All boxplots show median marked at center and data maxima and minima indicated by whiskers. Box shows 25th to 75th percentiles.

### *Xenopus* TPX2 NLS3 mutants increase recruitment of activated Aurora A and pericentrin to spindle poles

We next investigated which known protein interactions of TPX2 were altered in the presence of NLS3 mutants to mediate changes in spindle morphology and ectopic MT aster formation. TPX2 binds and allosterically maintains activation of Aurora A by protecting a phosphorylation site located in the activation loop at T295 in *Xenopus* (T288 in human) (Bayliss et al., 2003; Eyers & Maller, 2004). Using a phospho-specific antibody, we quantified total phospho-Aurora A T295 (pT295-AurA) immunostaining normalized to tubulin intensity on spindles generated in the presence of wildtype and NLS3 mutant *X. laevis* and *X. tropicalis* TPX2 proteins. Compared to GFP, addition of TPX2 proteins recruited and activated more Aurora A at spindle poles, with the *X. tropicalis* TPX2 NLS3 mutant showing the highest pT295-AurA staining, followed by wildtype *X. tropicalis*, *X. laevis* NLS3 mutant, and wildtype *X. laevis* (Figure 2A and B). Comparable differences in Aurora A activation were also detected biochemically by immunoprecipitating the recombinant proteins from egg extract reactions using a GFP antibody and probing Western blots with the pT295-AurA antibody (Figure 2C). To test whether the effects of the NLS3 mutant on MTs were mediated through kinase activation, we treated extracts with the Aurora A inhibitor Alisertib (MLN 8237), which was previously shown to diminish MTs emanating from sperm centrosomes and prevent spindle formation from Aurora A coated beads (Huang et al., 2018; Nadkarni & Heald, 2021). However, although MT density in spindles decreased with inhibitor treatment, pT295-AurA staining was not entirely abolished (Supplementary Figure 2), and NLS3 TPX2 mutants still co-precipitated with reduced levels of active kinase in the presence of the drug (Figure 2D). Together, these results show that mutating the NLS3 motif increases TPX2 binding and activation of Aurora A at spindle poles that is at least in part responsible for the increased MT polymerization observed.

**FIGURE 2:**
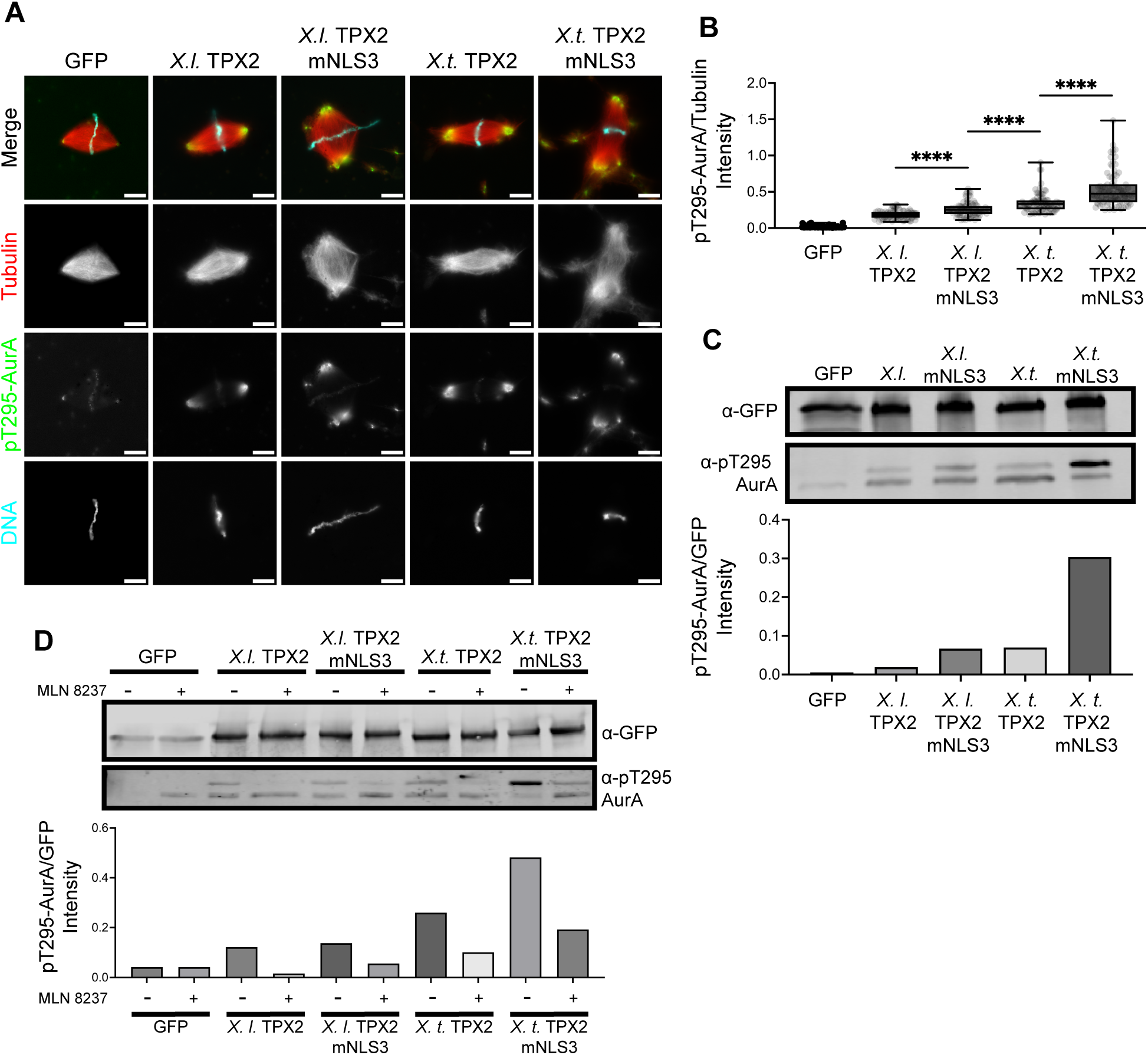
*X. laevis* and *X. tropicalis* TPX2 NLS3 motif mutants increase levels of pAurora A recruited to the spindle. (A) Immunostaining of spindle assembly reactions in *X. laevis* egg extracts in the presence of 100 nM recombinant GFP and GFP-tagged TPX2 proteins. Scale bars = 10 µm. (B) Ratio of fluorescence intensity of pT295-AurA/Tubulin on spindles, from three different extracts. n = 3 extracts, >27 spindles per replicate. Median marked at center and data maxima and minima indicated by whiskers. Box shows 25th to 75th percentiles. For each TPX2 protein compared with GFP control, **** = p<0.0001 from two-sided unpaired Welch’s *t* test. (C) Immunoprecipitation of GFP and GFP-TPX2 proteins from CSF aster reactions with 500 nM protein added followed by immunoblot analysis using indicated antibodies. (D) Immunoprecipitation of GFP and GFP-TPX2 proteins from CSF aster reactions in the presence of DMSO or 1 µM MLN 8237. 500 nM protein was added for each condition followed by immunoblot analysis using indicated antibodies.

In addition to inducing MT nucleation, Aurora A also regulates centrosome maturation and recruitment of pericentriolar material (PCM) (Magnaghi-Jaulin et al., 2019). To investigate whether TPX2 NLS3 mutants also affected PCM assembly, we examined localization of pericentrin (PCNT), a core PCM component (Doxsey et al., 1994). Quantification of total PCNT immunostaining intensity on spindles normalized to tubulin revealed that, consistent with effects on Aurora A activation, the *X. tropicalis* NLS3 mutant recruited the highest levels of pericentrin (Figure 3), which like several PCM scaffold proteins, has been shown to undergo phase separation in human cells (Jiang et al., 2021). In *C. elegans*, such condensates also contain XMAP215 as well as a TPX2-like protein (Woodruff et al., 2017). It is tempting to speculate that the hollow pole phenotype observed when MT growth is stimulated by TPX2, particularly the *X. tropicalis* NLS3 mutant, is due to the formation of concentrically layered condensates containing PCM as well as SAFs that recruit soluble tubulin and drive MT assembly at the periphery, similar to what has been observed in acentrosomal mouse oocyte spindles (Kraus, Alfaro-Aco, et al., 2023; So et al., 2019). Further experiments will need to examine the pole structures at higher resolution and localization of other PCM and SAF proteins.

**FIGURE 3:**
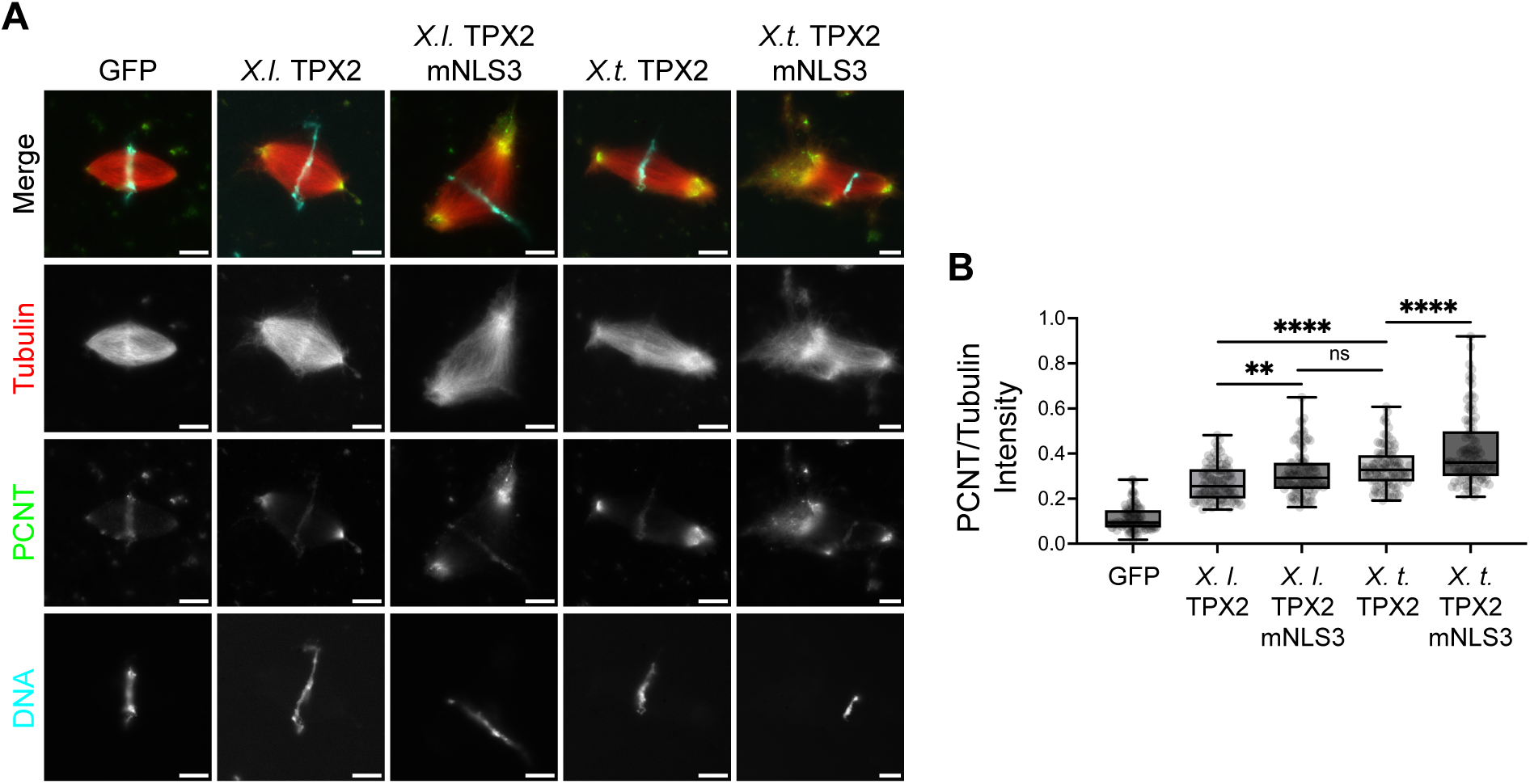
*X. laevis* and *X. tropicalis* TPX2 NLS3 motif mutants recruit more pericentrin (PCNT) to spindle poles. (A) Immunostaining of spindle assembly reactions in *X. laevis* egg extracts in the presence of 100 nM recombinant GFP or GFP-tagged TPX2 proteins. Scale bars = 10 µm. *X.t*. TPX2 mNLS3 image is cropped to show a larger field of view. (B) Ratio of fluorescence intensity of PCNT/Tubulin on spindles, from n = 3 extracts, >27 spindles per biological replicate. Median marked at center and data maxima and minima indicated by whiskers. Box shows 25th to 75th percentiles. For each TPX2 protein compared with GFP control, **** = p<0.0001 from two-sided unpaired Welch’s *t* test. ** = p<0.01, ns = not significant.

### *Xenopus* NLS3 TPX2 mutants increase recruitment of Eg5 and importin-α to spindle poles

An additional interacting protein of TPX2 is the class 5 kinesin-like motor protein Eg5 (Eckerdt et al., 2008). Previously we had shown that addition of recombinant TPX2 to *X. laevis* spindles increased the localization of Eg5 as well as MT bundling at spindle poles, altering spindle architecture (Helmke & Heald, 2014). This led us to test whether NLS3 mutants also affected Eg5 recruitment. Using immunostaining, we quantified total Eg5 intensity on spindles formed in the presence of each recombinant TPX2 protein, normalized to tubulin intensity, and observed that NLS3 mutations in both *X. laevis* and *X. tropicalis* increased Eg5 recruitment (Figure 4A). Interestingly, wildtype or mutant *X. tropicalis* TPX2 proteins recruited significantly more Eg5 to the spindle than the *X. laevis* proteins, indicating that *X. tropicalis* TPX2 binds more Eg5 (Figure 4B), which was corroborated by immunoprecipitation of the recombinant proteins from extract reactions and immunoblot analysis (Figure 4C). These findings indicate that the NLS3 motif acts additively to recruit Eg5 together with the previously identified deletion of a 7-a.a. sequence, 619-LSGSIVQ-625, in *X. tropicalis* TPX2. This 7-a.a. sequence is located just upstream of the C-terminal Eg5 binding domain (λ−7) (Helmke & Heald, 2014) and its deletion was shown to stimulate MT branching nucleation activity of *X. laevis* TPX2 (Alfaro-Aco et al., 2017). Mechanistically, it remains unknown how these two regions of TPX2 that are hundreds of amino acids apart modulate its activity.

**FIGURE 4:**
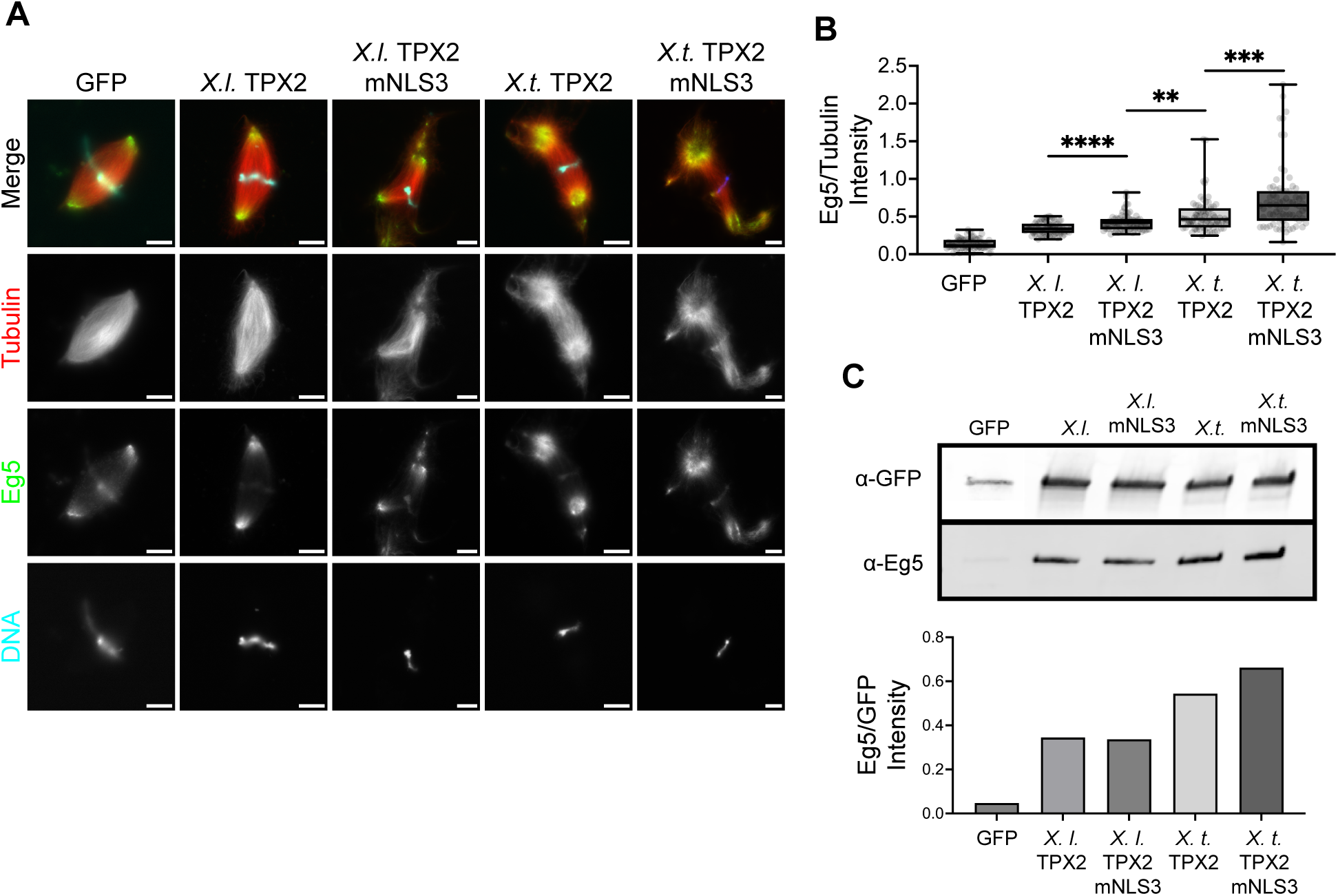
Increased recruitment of Eg5 to metaphase spindles upon addition of recombinant *X. tropicalis* wildtype and NLS3 mutant TPX2 proteins. (A) Immunostaining of spindle assembly reactions in *X. laevis* egg extracts in the presence of 100 nM recombinant GFP and GFP-tagged TPX2 proteins. Scale bars = 10 µm. *X.l.* TPX2 mNLS3 and *X.t*. TPX2 mNLS3 images are cropped to show a larger field of view. (B) Ratio of fluorescence intensity of Eg5/Tubulin on spindles, from n = 3 extracts, >23 spindles per biological replicate. Median marked at center and data maxima and minima indicated by whiskers. Box shows 25th to 75th percentiles. For each TPX2 protein compared with GFP control, **** = p<0.0001 from two-sided unpaired Welch’s *t* test. *** = p<0.001, ** = p<0.01. (C) Immunoprecipitation of GFP and GFP-TPX2 from CSF aster reactions supplemented with 500 nM recombinant protein followed by immunoblot analysis using indicated antibodies.

As a key Ran-regulated spindle assembly factor, TPX2 binds to importins and is released in the vicinity of mitotic chromosomes to stimulate MT polymerization and organization (Cavazza & Vernos, 2016; Gruss et al., 2001, 2002; Gruss & Vernos, 2004; Schatz et al., 2003; Walczak & Heald, 2008). Crystal structures of a truncated *X. laevis* TPX2 bound to importin-α demonstrated interaction through two NLS motifs located at 284-KRKH-287 (NLS1) and 327-KMIK-330 (NLS2) (Giesecke & Stewart, 2010). However, *in vitro* work measuring binding kinetics of TPX2 to importin-α showed that the NLS3 motif, 123-KKLK-126, alone confers a higher affinity for importin-α than the NLS1 and NLS2 sites do (Safari et al., 2021). We therefore investigated the contribution of TPX2 NLS3 to importin-α spindle localization and binding in egg extract. Immunofluorescence and quantification of total importin-α intensity on spindles normalized to tubulin revealed that addition of *X. laevis* NLS3 mutant TPX2 slightly reduced importin-α recruitment compared wildtype, while the *X. tropicalis* TPX2 NLS3 mutant showed a slight increase in importin-α levels (Figure 5A and B). By immunoprecipitation and immunoblot analysis, we found that both *X. laevis* and *X. tropicalis* TPX2 NLS3 mutant proteins still bound to importin-α (Figure 5C), which is perhaps not surprising since NLS1 and NLS2 were intact. The observation that *X. tropicalis* NLS3 mutant confers more importin-α binding to the spindle indicates again that the NLS3 motif acts additively with the Δ7.

**FIGURE 5:**
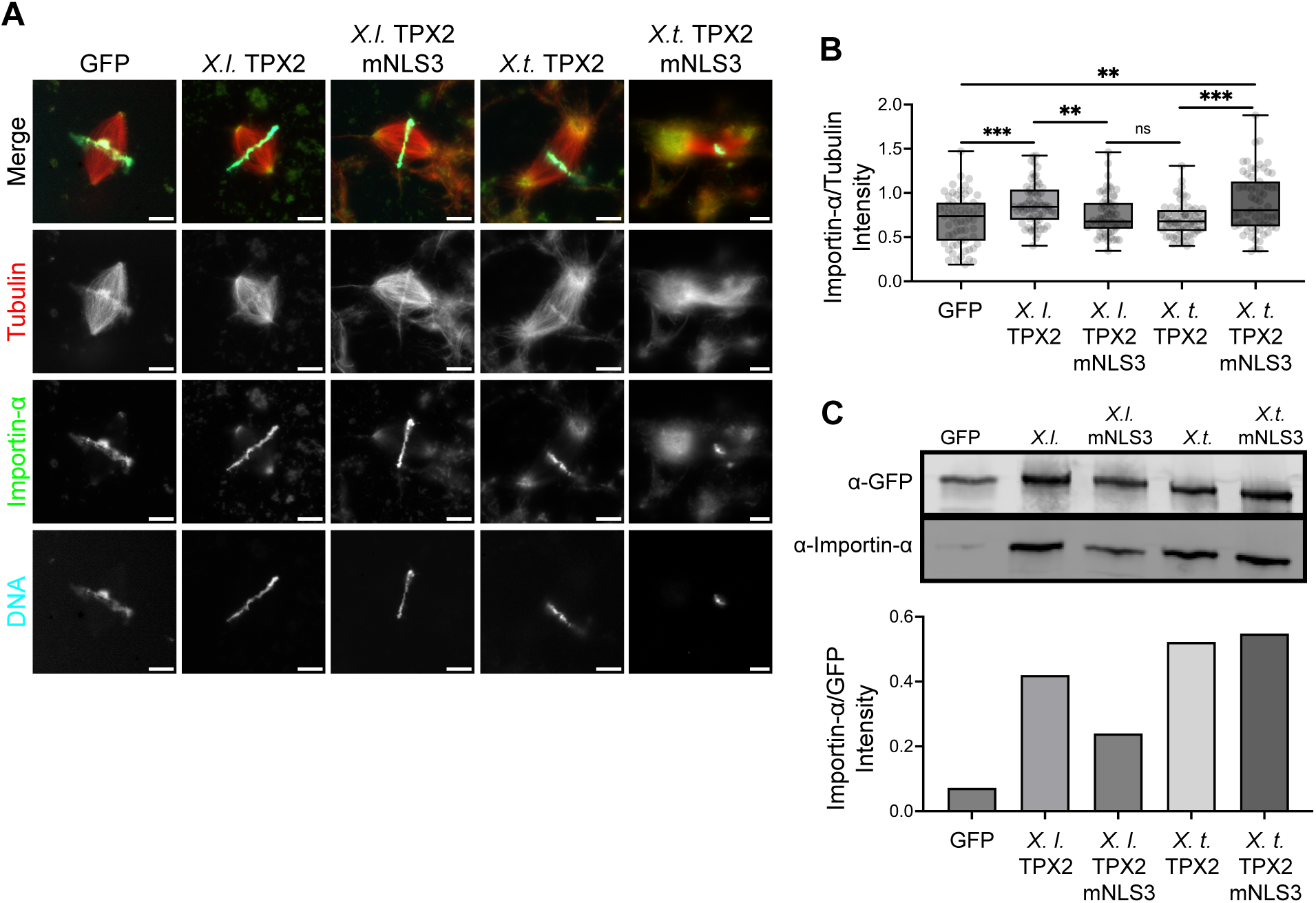
*X. laevis* and *X. tropicalis* TPX2 NLS3 mutants localize more importin-α to spindle poles. (A) Immunostaining of spindle assembly reactions in *X. laevis* egg extracts in the presence of 100 nm recombinant GFP and GFP-tagged TPX2 proteins. Scale bar = 10 µm. *X.t*. TPX2 mNLS3 image is cropped to show a larger field of view. (B) Ratio of fluorescence intensity of importin-α/tubulin on spindles from n = 3 extracts, >21, *** = p<0.001, ** = p<0.01 from two-sided Welch’s *t* test, ns = not significant. Median marked at center and data maxima and minima indicated by whiskers. Box shows 25th to 75th percentiles. (C) Immunoprecipitation of GFP and GFP-TPX2 from CSF aster reactions supplemented with 500 nM recombinant protein followed by immunoblot analysis using indicated antibodies.

Previous work has shown how importin-α can spatially regulate the minus-end-directed kinesin motor, XCTK2, on spindle MTs (Ems-McClung et al., 2004; Weaver et al., 2015). Near the chromatin, XCTK2 released from importins by RanGTP crosslinks MTs while also sliding both parallel and antiparallel MTs. At the spindle poles, XCTK2 MT crosslinking ability is selectively inhibited by importin-α, allowing for differential activity based on its location in the spindle (Ems-McClung et al., 2004; Weaver et al., 2015). In contrast, our results indicate that importin-α does not block TPX2 activity spindle poles.

Furthermore, increased recruitment of importin-α to spindle poles by *X. tropicalis* TPX2 NLS3 mutant may suggest an alternative role for importin-α at the spindle poles. *In vitro*, a truncated version of importin-α has been shown to enhance TPX2 condensation (Safari et al., 2021). There is also evidence that importin-α isoforms can form dimers *in vitro* and simultaneously bind to a single cargo containing a bipartite NLS (Matsuura, 2023; Miyamoto & Oka, 2016). Given these findings, importin-α could enhance TPX2 activity at the spindle poles by functioning as a point of multivalency, facilitating interactions among various SAFs. Future experiments involving TPX2 mutants lacking all NLSs will inform how importin-α recruitment contributes to LLPS, MT polymerization and spindle pole morphology.

Overall, our study demonstrates that an NLS mutation and a small deletion in TPX2 produce a gain of function phenotype by increasing recruitment of several TPX2-interacting proteins to spindle poles, increasing PCM levels, Aurora A activation, and astral MT polymerization. We also highlight how comparison of TPX2 from even closely related frog species provides insight into how differences in TPX2 sequence affect spindle architecture. Whether and how TPX2 activity is regulated at the transition between oocyte meiosis and zygotic mitosis, remains an important open question. Nonetheless, this work supports the notion that TPX2 is a linchpin for spindle assembly. Further exploring its structure-function relationships and regulation will be key to unraveling the role of TPX2 in determining spindle architecture and how changes in MT organization affect the fidelity of chromosome segregation.

## Material and Methods

### *Xenopus* egg extract and spindle assembly reactions

All frogs were used and maintained following standard protocols established by the UC Berkeley Animal Care and Use Committee and approved in our Animal Use Protocol. Egg extracts from *X. laevis* were prepared as previously described (Hannak & Heald, 2006; Maresca & Heald, 2006). Briefly, eggs were dejellied and then fractionated in a Sorvall HB-6 rotor for 16 min at 10,200 rpm. The cytoplasmic layer was then isolated and supplemented with 10 μg/ml LPC, 20 μg/ml CytoB, 1× energy mix, and 0.3 μM rhodamine-labeled porcine tubulin. Spindle reactions contained 50 μl cytostatic-factor (CSF) metaphase-arrested extract and *X. laevis* sperm at a final concentration of 500 nuclei per μl. Recombinant protein and inhibitors were added before the start of spindle assembly reactions, which were carried out in a 18°C water bath and flicked every 10– 15 min until spindle assembly was complete, approximately 45 min.

### Protein Purification

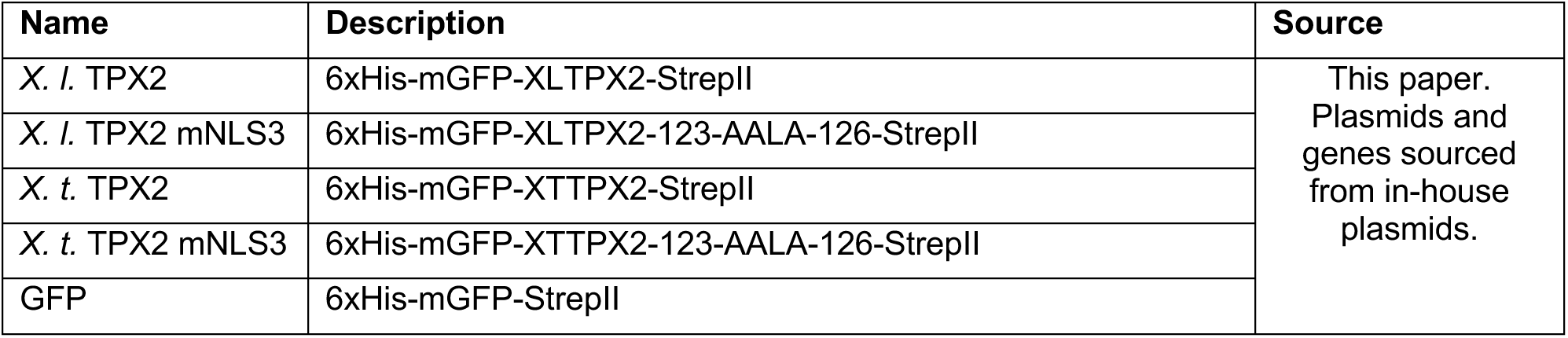

6xHis-mGFP-StrepII and 6xHis-mGFP-TPX2-StrepII *X. laevis* and *X. tropicalis* proteins were cloned into the pHAT2 vector (Addgene #112583) using NEBuilder HiFi DNA Assembly Master Mix (NEB #E2621). Proteins were transfected into Rosetta2 (DE3) pLysS *E. coli* cells (QB3 Berkeley Macro Lab) and grown to an OD of ∼0.35 at 37°C and then moved to 16°C until an OD of 0.5-0.7 was reached. Protein expression was induced with 750 nM Isopropyl ß-D-1-thiogalactopyranoside (IPTG) and carried out for 5-8 hours at 27°C. Cells were pelleted, resuspended in lysis buffer (50 mM Hepes, 20 mM Imidazole, 750 mM NaCl, 5 mM MgCl2, 25 mM sucrose, pH 7.75) containing 3 µM Avidin (Thermo Fisher # A887) 200 µM phenylmethylsulfonyl fluoride (PMSF), 10 μg/ml LPC, 5 mM β-mercaptoethanol (βME), cOmplete™ EDTA-free Protease Inhibitor tablet (Roche #11873580001), 10 µg/ml DNase I (Roche #10104159001) and flash frozen to be stored in −80°C for purification the following day. After thawing, resuspended cells were supplemented with Bug Buster protein reagent (Millipore #70584-3) before being sonicated. Lysate was centrifugated in a Sorvall SS-34 fixed angle rotor for 30 min at 4°C. Cleared lysate was flowed over pre-equilibrated HisPur™ Ni-NTA Resin (Thermo Fisher #88223) in a gravity column. The column was washed with 2x with 5 CV. Protein was eluted using lysis buffer with 0.5 M imidazole then underwent buffer exchange using via Sephadex G-25 in PD-10 Desalting Columns (Cytiva #17085101) into strep buffer (50 mM Hepes, 75 mM NaCl, 5 mM MgCl2, 25 mM sucrose, 5 mM EDTA, pH 7.75) containing 200 µM phenylmethylsulfonyl fluoride (PMSF), 10 μg/ml LPC, 5 mM β-mercaptoethanol (βME), cOmplete™ EDTA-free Protease Inhibitor tablet (Roche #11873580001). Eluant was then run over a pre-equilibrated Strep-Tactin^®^XT 4Flow^®^ high capacity resin (IBA # 2-5030-010) gravity column. The column was washed 3x with 2 column volumes and protein eluted using strep elution buffer (50 mM Hepes, 75 mM NaCl, 5 mM MgCl2, 50 mM arginine, 50 mM glutamate, 25 mM sucrose, 50 mM Biotin, pH 7.75). Eluant was concentrated using Amicon® Ultra Centrifugal Filter (Millipore #UFC9100) then buffer exchanged into XB buffer (10 mM Hepes, 2 mM MgCl2, 1 mM CaCl2, 10% w/v sucrose, pH 7.75) using Zeba spin desalting column (Thermo Fisher #87766).

### Immunofluorescence of spindles in egg extracts

Spindle reactions were fixed and processed for immunofluorescence as described in (Hannak & Heald, 2006). 50 µl reactions were fixed with a spindle dilution buffer (80 mM Pipes, 1 mM MgCl_2_, 1 mM EGTA, 30% glycerol, 0.5% Triton X-100) containing 3.7% formaldehyde for 5-10 min with rotation at 23°C. Fixed reaction were spun down onto coverslips through a cushion buffer (BRB80 + 40% glycerol) at 5,000 rpm for 20 min at 16°C using a Sorvall HS-4. Coverslips were postfixed for 5 min in 100% cold methanol, rinsed with PBS-0.1%NP40, blocked with PBS–3% BSA for 45 min. Primary antibodies against GFP (diluted 1:500), pT295-AurA (1: 1,000), PCNT (1:250), Eg5 (1: 2,500), or importin-α (1: 2000) were added overnight in a humidified chamber at 4°C. The following steps were carried out at room temperature. Coverslips were rinsed 3x quickly with PBS-0.1%NP40, then 3x for 5 min. Secondary antibodies (Invitrogen; Goat anti-rabbit or Goat anti-rabbit IGG conjugated to Alexa Fluor 488, 1:500) were added and incubated for 30 min. Coverslips were rinsed 3x quickly with PBS-0.1%NP40, then 3x for 5 min. 5 μg/ml Hoechst 33258 (Sigma) was added for 10 min before mounting with ProLong™ Glass Antifade Mountant (Thermo Fisher).

### Imaging and Analysis

Images were acquired using an epifluorescence Olympus BX51 microscope with an Olympus PlanApo 63x oil objective and Hamamatsu ORCA-II camera. Exposure times for each channel were set for the sample with the highest fluorescence intensity and kept constants for all replicates of that experiment. Images were cropped using a consistent area and all fluorescence intensities were measure using FIJI software. Statistical significance was determined by unpaired two sample Welch’s *t* test using GraphPad Prism version 10.0.0 for MacOS, GraphPad Software, Boston, Massachusetts, USA, www.graphpad.com. p values are listed in the figure legend.

### Immunoprecipitation and Western blotting

CSF aster reactions were used to perform all immunoprecipitation experiments. Reactions were set up similarly to spindle reactions except without sperm DNA and in the presence of 500 nM recombinant protein for each condition. Proteins and 1µM MLN 8237 inhibitor or DMSO were added prior to initiation of extract reactions, which were incubated for 40 min in an 18°C water bath and flicked every 10–15 min. Each reaction was then added to Strep-Tactin^®^XT 4Flow^®^ high capacity resin (IBA # 2-5030-010) pre-equilibrated with XB buffer supplemented with cOmplete™ EDTA-free Protease Inhibitor tablets (Roche #11873580001) and/or PhosSTOP™ Phosphatase Inhibitor Tablets (Roche # 04906845001). Samples incubated with rotation at 4°C were each washed with 700 µl of XB buffer. After the initial wash, samples were pelleted at max speed for 1 min in a tabletop centrifuge. Supernatant was carefully removed followed by two more washes with 700 µl XB buffer + 0.1% triton x-100. After the last wash, samples were pelleted and resuspended in 1X SDS sample buffer and boiled for 10 min. For Western blot analysis, 25% of sample was run on a SDS-PAGE gel. Wet transferers were carried out using nitrocellulose membrane (Thermo Fisher #). Blots were blocked with either 5% milk + PBS-0.1% Tween or 5% BSA + PBS-0.1% Tween for 1 hour at room temperature then washed with 1xTBST or 1xTBS 3x quickly then 3x for 5 min. Primary antibodies diluted in blocking buffer against GFP (1:1000), pT295-AurA (1:1,000), Eg5 (1:2,500), or importin-α (1:2000) were incubated overnight at 4°C with rotation. Blots were washed with 1xTBST or 1xTBS 3x quickly then 3x for 5 min before incubating with secondary antibodies Alexa Fluor 700 goat-anti-rabbit IgG (Invitrogen # A21038) or IRDye-800CW goat anti-mouse IgG (LI-COR # 926-32210) diluted in blocking buffer for 1 hour at room temperature. Blots were washed with 1xTBST or 1xTBS 3x quickly then 3x for 5 min then scanned on an Odyssey Infrared Imaging System (LI-COR Biosciences). Band intensities were quantified using Fiji.

## Acknowledgements

We thank Taron Kapoor for his generous donation of Eg5 antibody. We also thank Coral Zhou and members of the Heald lab for assistance in data analysis and comments on the manuscript. This work was funded by NIH MIRA grant R35GM118183 and the Flora Lamson Hewlett Chair in Biochemistry to R.H. and a Cancer Research Coordinating Committee Fellowship to G.S.

---

**SUPPLEMENTARY FIGURE 1:**
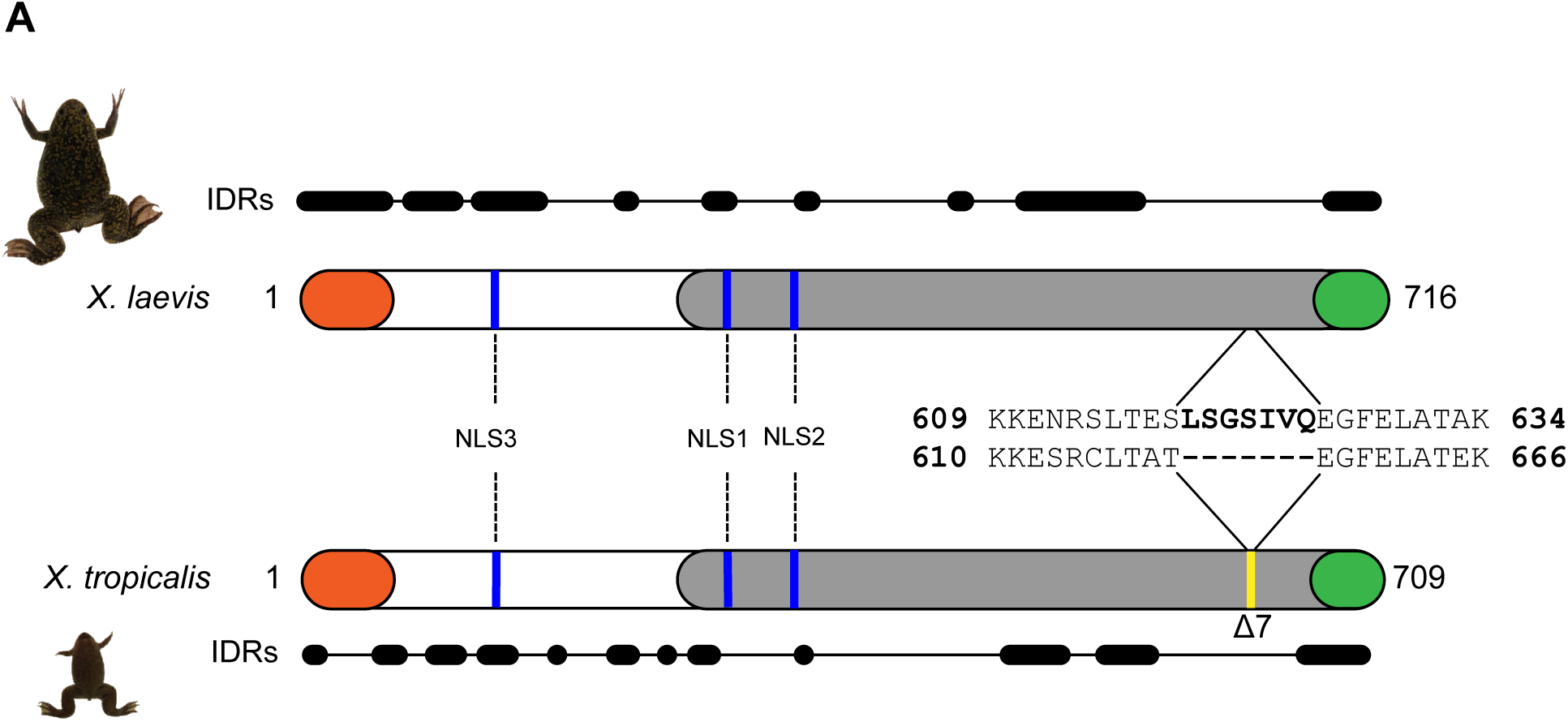
Schematic of TPX2 domains in *X. laevis* and *X. tropicalis*. N-terminal Aurora A binding domain (orange; (Bayliss et al., 2003)) and C-terminal Eg5 binding domain (green; (Bayliss et al., 2003; Eckerdt et al., 2008)). Mapped NLS1 (^284^KRKH^287^), NLS2 (^327^KMIK^330^) (Giesecke & Stewart, 2010) and NLS3 (^123^KKLK^126^) (Safari et al., 2021) *X. tropicalis* NLS2 (^328^KMVR^331^). Region known to interact with MT and stimulate nucleation *in vitro* (gray; (Roostalu et al., 2015; Zhang et al., 2017)). Zoomed in view of sequence alignment between *X. laevis* and *X. tropicalis* highlighting 7-aa deletion (Δ7) in *X. tropicalis* (yellow; (Helmke & Heald, 2014)). Disordered regions predicted using fIDPnn program (Hu et al., 2021).

**SUPPLEMENTARY FIGURE 2:**
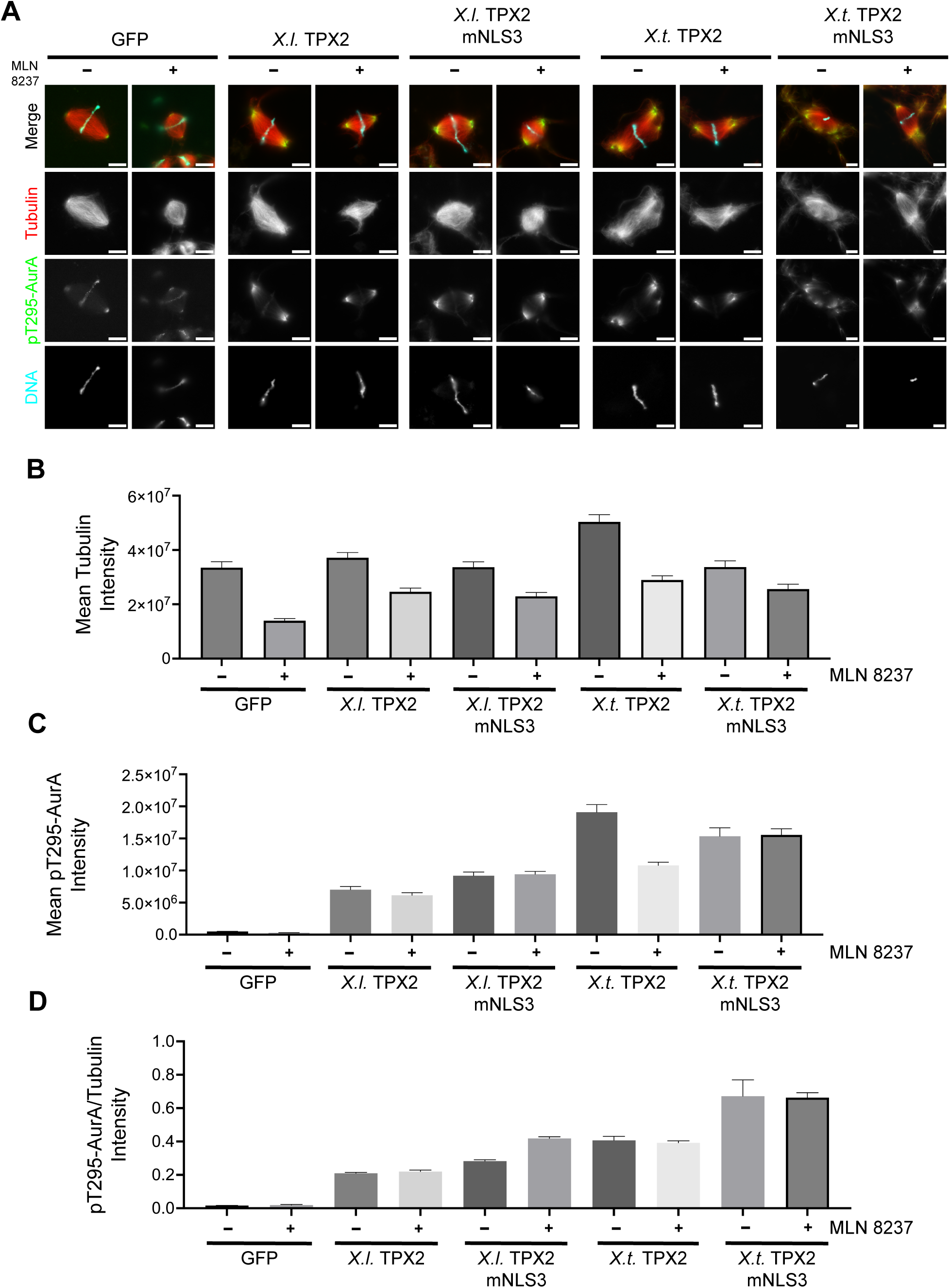
Inhibition of Aurora A by MLN 8237 inhibitor decreases tubulin intensity in spindles in the presence of *X. laevis* and *X. tropicalis* TPX2 NLS3 motif mutants. (A) Immunostaining of spindle assembly reactions in *X. laevis* egg extracts with 100 nM recombinant GFP and GFP-tagged TPX2 proteins and DMSO or 1 µM MLN 8237. Scale bars = 10 µm. *X.t*. TPX2 mNLS3 image is cropped to show a larger field of view. (B) Mean fluorescence intensity of tubulin, (C) Mean fluorescence intensity of p295T-Aurora A, and (D) Ratio of fluorescence intensity of pT295-AurA/Tubulin on spindles, n = 3 biological replicates, >25 spindles per replicate. Error bars = SEM.

## References

Alfaro-Aco, R., Thawani, A., & Petry, S. (2017). Structural analysis of the role of TPX2 in branching microtubule nucleation. Journal of Cell Biology, 216(4), 983–997. 10.1083/jcb.201607060

Asteriti, I. A., Rensen, W. M., Lindon, C., Lavia, P., & Guarguaglini, G. (2010). The Aurora-A/TPX2 complex: A novel oncogenic holoenzyme? Biochimica et Biophysica Acta (BBA) - Reviews on Cancer, 1806(2), 230–239. 10.1016/j.bbcan.2010.08.001

Bayliss, R., Sardon, T., Vernos, I., & Conti, E. (2003). Structural Basis of Aurora-A Activation by TPX2 at the Mitotic Spindle. Molecular Cell, 12(4), 851–862. 10.1016/s1097-2765(03)00392-7

Brunet, S., Sardon, T., Zimmerman, T., Wittmann, T., Pepperkok, R., Karsenti, E., & Vernos, I. (2004). Characterization of the TPX2 Domains Involved in Microtubule Nucleation and Spindle Assembly in Xenopus Egg Extracts. Molecular Biology of the Cell, 15(12), 5318–5328. 10.1091/mbc.e04-05-0385

Cavazza, T., & Vernos, I. (2016). The RanGTP Pathway: From Nucleo-Cytoplasmic Transport to Spindle Assembly and Beyond. Frontiers in Cell and Developmental Biology, 3, 82. 10.3389/fcell.2015.00082

Crowder, M. E., Strzelecka, M., Wilbur, J. D., Good, M. C., von Dassow, G., & Heald, R. (2015). A Comparative Analysis of Spindle Morphometrics across Metazoans. Current Biology, 25(11), 1542–1550. 10.1016/j.cub.2015.04.036

Doxsey, S. J., Stein, P., Evans, L., Calarco, P. D., & Kirschner, M. (1994). Pericentrin, a highly conserved centrosome protein involved in microtubule organization. Cell, 76(4), 639–650. 10.1016/0092-8674(94)90504-5

Eckerdt, F., Eyers, P. A., Lewellyn, A. L., Prigent, C., & Maller, J. L. (2008). Spindle Pole Regulation by a Discrete Eg5-Interacting Domain in TPX2. Current Biology, 18(7), 519–525. 10.1016/j.cub.2008.02.077

Ems-McClung, S. C., Zheng, Y., & Walczak, C. E. (2004). Importin α/β and Ran-GTP Regulate XCTK2 Microtubule Binding through a Bipartite Nuclear Localization Signal. Molecular Biology of the Cell, 15(1), 46–57. 10.1091/mbc.e03-07-0454

Eyers, P. A., Erikson, E., Chen, L. G., & Maller, J. L. (2003). A Novel Mechanism for Activation of the Protein Kinase Aurora A. Current Biology, 13(8), 691–697. 10.1016/s0960-9822(03)00166-0

Eyers, P. A., & Maller, J. L. (2004). Regulation of Xenopus Aurora A Activation by TPX2*. Journal of Biological Chemistry, 279(10), 9008–9015. 10.1074/jbc.m312424200

Gable, A., Qiu, M., Titus, J., Balchand, S., Ferenz, N. P., Ma, N., Collins, E. S., Fagerstrom, C., Ross, J. L., Yang, G., & Wadsworth, P. (2012). Dynamic reorganization of Eg5 in the mammalian spindle throughout mitosis requires dynein and TPX2. Molecular Biology of the Cell, 23(7), 1254–1266. 10.1091/mbc.e11-09-0820

Garrett, S., Auer, K., Compton, D. A., & Kapoor, T. M. (2002). hTPX2 Is Required for Normal Spindle Morphology and Centrosome Integrity during Vertebrate Cell Division. Current Biology, 12(23), 2055–2059. 10.1016/s0960-9822(02)01277-0

Giesecke, A., & Stewart, M. (2010). Novel Binding of the Mitotic Regulator TPX2 (Target Protein for Xenopus Kinesin-like Protein 2) to Importin-α*. Journal of Biological Chemistry, 285(23), 17628–17635. 10.1074/jbc.m110.102343

Gruss, O. J., Carazo-Salas, R. E., Schatz, C. A., Guarguaglini, G., Kast, J., Wilm, M., Bot, N. L., Vernos, I., Karsenti, E., & Mattaj, I. W. (2001). Ran Induces Spindle Assembly by Reversing the Inhibitory Effect of Importin α on TPX2 Activity. Cell, 104(1), 83–93. 10.1016/s0092-8674(01)00193-3

Gruss, O. J., & Vernos, I. (2004). The mechanism of spindle assembly. The Journal of Cell Biology, 166(7), 949–955. 10.1083/jcb.200312112

Gruss, O. J., Wittmann, M., Yokoyama, H., Pepperkok, R., Kufer, T., Silljé, H., Karsenti, E., Mattaj, I. W., & Vernos, I. (2002). Chromosome-induced microtubule assembly mediated by TPX2 is required for spindle formation in HeLa cells. Nature Cell Biology, 4(11), 871–879. 10.1038/ncb870

Hannak, E., & Heald, R. (2006). Investigating mitotic spindle assembly and function in vitro using Xenopus laevis egg extracts. Nature Protocols, 1(5), 2305–2314. 10.1038/nprot.2006.396

Helmke, K. J., & Heald, R. (2014). TPX2 levels modulate meiotic spindle size and architecture in Xenopus egg extracts. Journal of Cell Biology, 206(3), 385–393. 10.1083/jcb.201401014

Hu, G., Katuwawala, A., Wang, K., Wu, Z., Ghadermarzi, S., Gao, J., & Kurgan, L. (2021). flDPnn: Accurate intrinsic disorder prediction with putative propensities of disorder functions. Nature Communications, 12(1), 4438. 10.1038/s41467-021-24773-7

Huang, Y., Li, T., Ems-McClung, S. C., Walczak, C. E., Prigent, C., Zhu, X., Zhang, X., & Zheng, Y. (2018). Aurora A activation in mitosis promoted by BuGZ. Journal of Cell Biology, 217(1), 107–116. 10.1083/jcb.201706103

Jiang, X., Ho, D. B. T., Mahe, K., Mia, J., Sepulveda, G., Antkowiak, M., Jiang, L., Yamada, S., & Jao, L.-E. (2021). Condensation of pericentrin proteins in human cells illuminates phase separation in centrosome assembly. Journal of Cell Science, 134(14), jcs258897. 10.1242/jcs.258897

King, M. R., & Petry, S. (2020). Phase separation of TPX2 enhances and spatially coordinates microtubule nucleation. Nature Communications, 11(1), 270. 10.1038/s41467-019-14087-0

Kraus, J., Alfaro-Aco, R., Gouveia, B., & Petry, S. (2023). Microtubule nucleation for spindle assembly: one molecule at a time. Trends in Biochemical Sciences, 48(9), 761–775. 10.1016/j.tibs.2023.06.004

Kraus, J., Travis, S. M., King, M. R., & Petry, S. (2023). Augmin is a Ran-regulated spindle assembly factor. Journal of Biological Chemistry, 299(6), 104736. 10.1016/j.jbc.2023.104736

Kufer, T. A., Silljé, H. H. W., Körner, R., Gruss, O. J., Meraldi, P., & Nigg, E. A. (2002). Human TPX2 is required for targeting Aurora-A kinase to the spindle. The Journal of Cell Biology, 158(4), 617–623. 10.1083/jcb.200204155

Ma, N., Titus, J., Gable, A., Ross, J. L., & Wadsworth, P. (2011). TPX2 regulates the localization and activity of Eg5 in the mammalian mitotic spindle. Journal of Cell Biology, 195(1), 87–98. 10.1083/jcb.201106149

Ma, N., Tulu, U. S., Ferenz, N. P., Fagerstrom, C., Wilde, A., & Wadsworth, P. (2010). Poleward Transport of TPX2 in the Mammalian Mitotic Spindle Requires Dynein, Eg5, and Microtubule Flux. Molecular Biology of the Cell, 21(6), 979–988. 10.1091/mbc.e09-07-0601

Magnaghi-Jaulin, L., Eot-Houllier, G., Gallaud, E., & Giet, R. (2019). Aurora A Protein Kinase: To the Centrosome and Beyond. Biomolecules, 9(1), 28. 10.3390/biom9010028

Maresca, T. J., & Heald, R. (2006). Xenopus Protocols, Cell Biology and Signal Transduction. Methods in Molecular Biology^TM^, 322, 459–474. 10.1007/978-1-59745-000-3_33

Matsuura, Y. (2023). Crystallographic data of an importin-α3 dimer in which the two protomers are bridged by a bipartite nuclear localization signal. Data in Brief, 47, 108988. 10.1016/j.dib.2023.108988

Miyamoto, Y., & Oka, M. (2016). Data on dimer formation between importin α subtypes. Data in Brief, 7, 1248–1253. 10.1016/j.dib.2016.03.080

Nadkarni, A. V., & Heald, R. (2021). Reconstitution of muscle cell microtubule organization in vitro. Cytoskeleton, 78(10–12), 492–502. 10.1002/cm.21710

Neumayer, G., Belzil, C., Gruss, O. J., & Nguyen, M. D. (2014). TPX2: of spindle assembly, DNA damage response, and cancer. Cellular and Molecular Life Sciences, 71(16), 3027–3047. 10.1007/s00018-014-1582-7

Petry, S., Groen, A. C., Ishihara, K., Mitchison, T. J., & Vale, R. D. (2013). Branching Microtubule Nucleation in Xenopus Egg Extracts Mediated by Augmin and TPX2. Cell, 152(4), 768–777. 10.1016/j.cell.2012.12.044

Pinyol, R., Scrofani, J., & Vernos, I. (2013). The Role of NEDD1 Phosphorylation by Aurora A in Chromosomal Microtubule Nucleation and Spindle Function. Current Biology, 23(2), 143–149. 10.1016/j.cub.2012.11.046

Roostalu, J., Cade, N. I., & Surrey, T. (2015). Complementary activities of TPX2 and chTOG constitute an efficient importin-regulated microtubule nucleation module. Nature Cell Biology, 17(11), 1422–1434. 10.1038/ncb3241

Safari, M. S., King, M. R., Brangwynne, C. P., & Petry, S. (2021). Interaction of spindle assembly factor TPX2 with importins-α/β inhibits protein phase separation. Journal of Biological Chemistry, 297(3), 100998. 10.1016/j.jbc.2021.100998

Sardon, T., Peset, I., Petrova, B., & Vernos, I. (2008). Dissecting the role of Aurora A during spindle assembly. The EMBO Journal, 27(19), 2567–2579. 10.1038/emboj.2008.173

Schatz, C. A., Santarella, R., Hoenger, A., Karsenti, E., Mattaj, I. W., Gruss, O. J., & Carazo-Salas, R. E. (2003). Importin α-regulated nucleation of microtubules by TPX2. The EMBO Journal, 22(9), 2060–2070. 10.1093/emboj/cdg195

Scrofani, J., Sardon, T., Meunier, S., & Vernos, I. (2015). Microtubule Nucleation in Mitosis by a RanGTP-Dependent Protein Complex. Current Biology, 25(2), 131–140. 10.1016/j.cub.2014.11.025

So, C., Seres, K. B., Steyer, A. M., Mönnich, E., Clift, D., Pejkovska, A., Möbius, W., & Schuh, M. (2019). A liquid-like spindle domain promotes acentrosomal spindle assembly in mammalian oocytes. Science, 364(6447). 10.1126/science.aat9557

Song, J.-G., King, M. R., Zhang, R., Kadzik, R. S., Thawani, A., & Petry, S. (2018). Mechanism of how augmin directly targets the γ-tubulin ring complex to microtubules. Journal of Cell Biology, 217(7), 2417–2428. 10.1083/jcb.201711090

Thawani, A., Kadzik, R. S., & Petry, S. (2018). XMAP215 is a microtubule nucleation factor that functions synergistically with the γ-tubulin ring complex. Nature Cell Biology, 20(5), 575–585. 10.1038/s41556-018-0091-6

Travis, S. M., Mahon, B. P., & Petry, S. (2022). How Microtubules Build the Spindle Branch by Branch. Annual Review of Cell and Developmental Biology, 38(1), 1–23. 10.1146/annurev-cellbio-120420-114559

Tsai, M.-Y., Wiese, C., Cao, K., Martin, O., Donovan, P., Ruderman, J., Prigent, C., & Zheng, Y. (2003). A Ran signalling pathway mediated by the mitotic kinase Aurora A in spindle assembly. Nature Cell Biology, 5(3), 242–248. 10.1038/ncb936

Tulu, U. S., Fagerstrom, C., Ferenz, N. P., & Wadsworth, P. (2006). Molecular Requirements for Kinetochore-Associated Microtubule Formation in Mammalian Cells. Current Biology, 16(5), 536–541. 10.1016/j.cub.2006.01.060

Walczak, C. E., & Heald, R. (2008). Chapter Three Mechanisms of Mitotic Spindle Assembly and Function. International Review of Cytology, 265, 111–158. 10.1016/s0074-7696(07)65003-7

Weaver, L. N., Ems-McClung, S. C., Chen, S.-H. R., Yang, G., Shaw, S. L., & Walczak, C. E. (2015). The Ran-GTP Gradient Spatially Regulates XCTK2 in the Spindle. Current Biology, 25(11), 1509–1514. 10.1016/j.cub.2015.04.015

Wittmann, T., Wilm, M., Karsenti, E., & Vernos, I. (2000). Tpx2, a Novel Xenopus Map Involved in Spindle Pole Organization. The Journal of Cell Biology, 149(7), 1405–1418. 10.1083/jcb.149.7.1405

Woodruff, J. B., Gomes, B. F., Widlund, P. O., Mahamid, J., Honigmann, A., & Hyman, A. A. (2017). The Centrosome Is a Selective Condensate that Nucleates Microtubules by Concentrating Tubulin. Cell, 169(6), 1066–1077.e10. 10.1016/j.cell.2017.05.028

Zhang, R., Roostalu, J., Surrey, T., & Nogales, E. (2017). Structural insight into TPX2-stimulated microtubule assembly. ELife, 6, e30959. 10.7554/elife.30959

